# Tissue resident macrophages innately develop in a human iPSC-derived cardiac organoid model

**DOI:** 10.1101/2025.05.20.654824

**Authors:** Anna Rockel, Tobias Brunnbauer, Nicole Wagner, Brenda Gerull, Süleyman Ergün, Philipp Wörsdörfer

**Affiliations:** Institute of Anatomy and Cell Biology, University of Würzburg, Germany; Comprehensive Heart Failure Center (CHFC) and Department of Internal Medicine I, University Hospital Würzburg, Würzburg, Germany

## Abstract

The heart is the first functional organ to develop during embryogenesis, forming in parallel with the vasculature and hematopoietic cell lineages. To advance our understanding of human cardiac development and disease, human induced pluripotent stem cell-derived cardiomyocytes offer a promising *in vitro* model. However, conventional 2D culture systems lack the complexity required to recapitulate the intricate interactions of different cell types leading to fully functional and mature cardiac tissue. Here, we present a cardiac organoid model that mimics several aspects of cardiogenesis. The organoids develop a functional myocardium consisting of cardiomyocytes and fibroblasts capable of spontaneous rhythmic contractions. The myocard is interspersed with a branched endothelial network. Additionally, macrophages develop within the organoids and integrate into the myocardium. In summary, we describe a complex 3D cell culture platform to study human heart tissue development with all the involved cell types (cardiomyocytes, fibroblasts, endothelial cells, macrophages), paving the way for new insights into the role of macrophages in cardiac development and disease.

## Introduction

In recent years, several human induced pluripotent stem cell (iPSC)-derived self-organizing cardiac organoid models were developed induced by biphasic modulation of the WNT-signaling pathway (Lewis-Israeli et al., 2021) (Hofbauer et al., 2021) (Drakhlis et al., 2021) (Volmert et al., 2023). These models generate different cell types of the heart, such as cardiomyocytes, fibroblasts, endothelial cells, epicardial cells, and additional non-cardiac tissues like foregut epithelium. However, the presence of tissue-resident macrophages has not been described yet.

Macrophages are a crucial component of the heart *in vivo*. They play a role in cardiac development (Xu et al., 2024), maintain cardiac tissue homeostasis (Nicolas-Avila et al., 2020), support electric conduction (Hulsmans et al., 2017), and contribute to the repair and regeneration of cardiac tissue (Zaman and Epelman, 2022). Moreover, recent studies have demonstrated improved contractile force, tissue functionality, and long-term vascularization when macrophages are incorporated into engineered heart tissues (Landau et al., 2024) (Lock et al., 2024) (Hamidzada et al., 2024).

Therefore, macrophages, included into cardiac organoids, would create a more physiologic model that better mimics the complex interactions within the heart.

Here, we describe the innate development of macrophages in cardiac organoids, an aspect that, to our knowledge, has not been reported before. Cardiac organoid macrophages develop from hemogenic endothelium within blood vessel-like structures and integrate into the cardiac muscle. They display tissue-resident macrophage-like characteristics as determined by single-cell RNA sequencing. In addition to macrophages, erythrocytes also develop indicating primitive hematopoiesis.

In summary, the described cardiac organoid model offers an advanced platform for studying human cardiogenesis, vascular development, and the involvement of macrophages, in these processes. Unlike previous models that often rely on exogenous addition of macrophages or lack them entirely, this system supports the innate emergence of macrophages. This adds a new layer of physiological relevance and complexity.

## Results

To generate the cardiac organoids, modifications were made to a previously published protocol for self-assembling cardioids (Lewis-Israeli et al., 2021). In contrast to the published method, the WNT antagonist IWP2 was used instead of WNT-C59, and the organoids were cultured on an orbital shaker. Of note, IWP2 was used at a low concentration of 2 µM. Finally, the second supplementation with CHIR99021, which was used in the original protocol to induce epicardial cells, was omitted (Fig. 1A).

**Figure 1.**
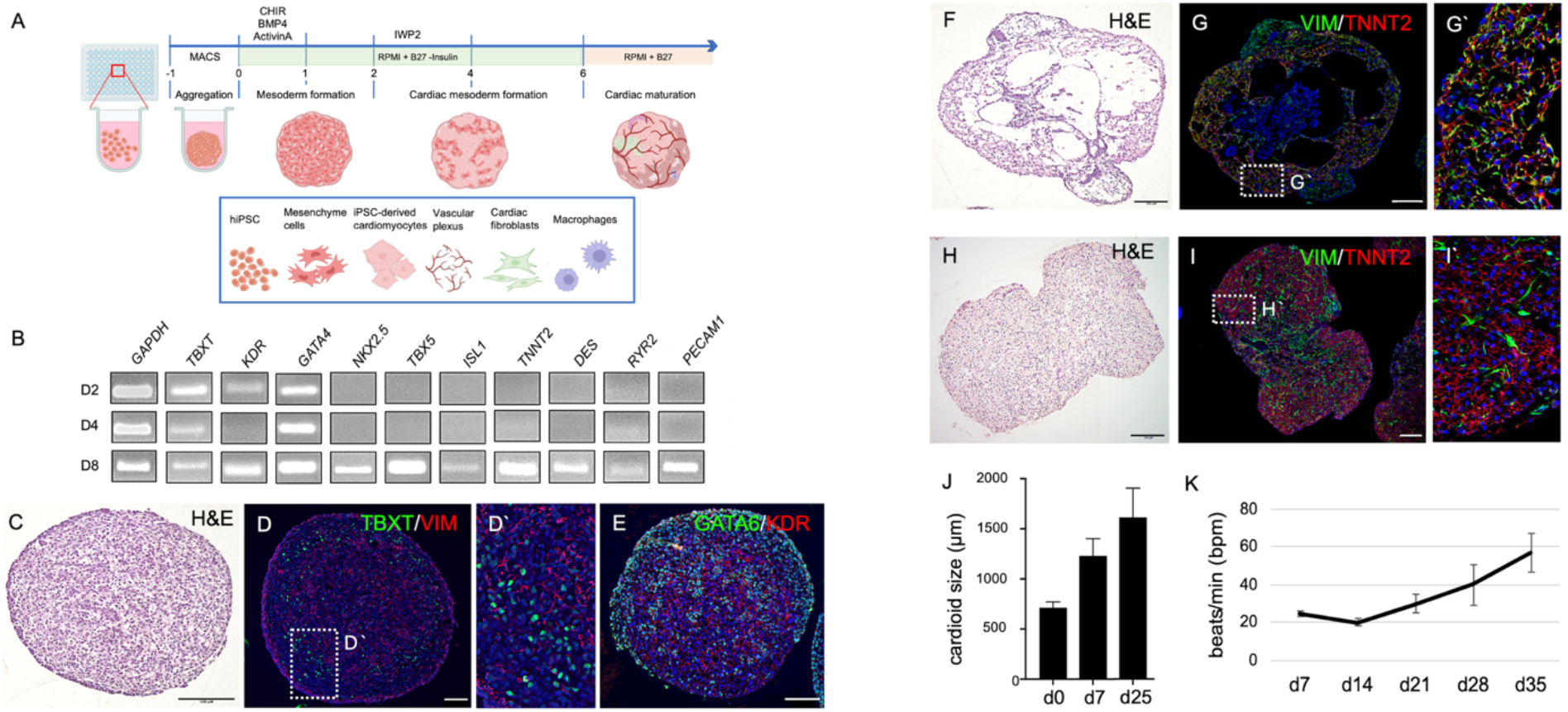
Cardiac Organoids. **A)** Schematic representation of the cardiac organoid differentiation protocol. **B)** Semiquantitative RT-PCR detecting mesodermal (*TBXT, KDR, GATA4*), cardiac (*NKX2*.*5, TBX5, ISL1, TNNT2, DES, RYR2*), and endothelial (*PECAM1*) marker gene expression. **C)** H&E staining of organoid section at day 2. **D-D’)** Immunofluorescence analyses detecting early mesodermal markers TBXT and VIM. **E)** Immunofluorescence analyses detecting early mesodermal markers GATA6 and KDR. **F)** H&E staining showing hollow cavity-like structures in cardiac organoids at day 8. **G-G’)** Immunofluorescence analyses detecting mesenchymal cells (VIM^+^) in close association with cardiomyocytes (TNNT2^+^). **H)** H&E staining of organoid sections at day 35 of culture. **I)** Immunofluorescence analyses showing close interaction of fibroblasts (VIM^+^) and cardiomyocytes (TNNT2^+^) at day 35 of culture. **J)** Cardiac organoids progressively gain in size (D0 = 12.2 μm ± 6.85; D7 = 1231 ± 20.31; D25 = 1613 ± 45.43). **K)** Beating frequency increases over culture time starting from 25 bpm (± 1.9) at day 7 up to 57 bpm (± 10.5) at day 35. Scale in all pictures: 50 µm

Initial characterizations using semiquantitative RT-PCR, demonstrate that the expression of marker genes for mesoderm specification (*TBXT, KDR, GATA4*), heart development (*NKX2*.*5, TBX5, ISL1*), cardiomyocytes (*TNNT2, DES, RYR2*), and endothelial cells (*PECAM1*) becomes upregulated over the course of differentiation. While mesodermal markers were detectable at day 2, cardiac and vascular transcripts were not detected before day 8 (Fig. 1B).

Histological analyses confirm these results (Fig. 1C). Immunofluorescence staining on paraffin sections of day 2 organoids demonstrate the expression of the mesodermal transcription factor TBXT in several nuclei (Fig. 1D). Moreover, expression of the mesenchymal marker Vimentin (VIM) is detectable (Fig. 1D). Furthermore, we observe GATA6 expression and expression of the VEGF receptor 2 (KDR), typical hallmarks of mesoderm formation (Fig. 1E). In day 8 organoids (Fig. 1F), the tissue is loosely arranged with large cavities and a core of epithelial cells in the center. The mesenchymal/fibroblast marker VIM is expressed in cells in close association with cells expressing the cardiomyocyte marker TNNT2. Both cell types are located in the periphery of the organoid (Fig. 1F-G). At day 35, the tissue architecture of the organoid becomes more compact, and initially formed cavities are lost (Fig. 1H). The organoid now consists mostly of cardiac tissue formed by VIM^+^ fibroblasts and TNNT2^+^ cardiomyocytes (Fig. 1I). During differentiation, organoids gradually increase in size from a diameter of approximately 700 µm (712.2 µm ± 6.8) at day 0 to a diameter of 1.6 mm at day 25 (1600 µm ± 45) (Fig. 1J). Spontaneous contractions start between day 7 and day 10, and beating frequencies increase from initially 25 bpm (±1.9) to approximately 60 bpm (57 bpm ± 10.5) after 4 weeks in culture (Fig. 1K).

In day 8 organoids, ACTN1 and TNNT2 exhibit a diffuse staining pattern, with only a few immature, sarcomere-like structures visible. By contrast, at day 35, a continuous cross-striated staining pattern emerges, indicating a more organized actin-myosin arrangement within the sarcomeres. This progression reflects the functional maturation of cardiac muscle over time. (Fig. 2A-B). Electron microscopic analyses confirmed the formation of well-organized sarcomeres with clearly visible Z-lines, as well as A-and I-bands and the H-zone. Sarcomeres at day 35 show a length of approximately 1.9 µm (1.89 µm ± 0.2; n=26). In addition, intercalated disc-like structures are observed between neighboring cardiomyocytes (Fig. 2F). Multiple mitochondria are found near the sarcomeres (Fig. 2C-D). The stimulation with 100 µM isoproterenol clearly increases the beating frequency (untreated: 18 bpm (±4.5), treated: 30 bpm (±5.3); n=9; P=<0.0001), demonstrating a response of the cardiac tissue to beta-adrenergic stimulation.

**Figure 2.**
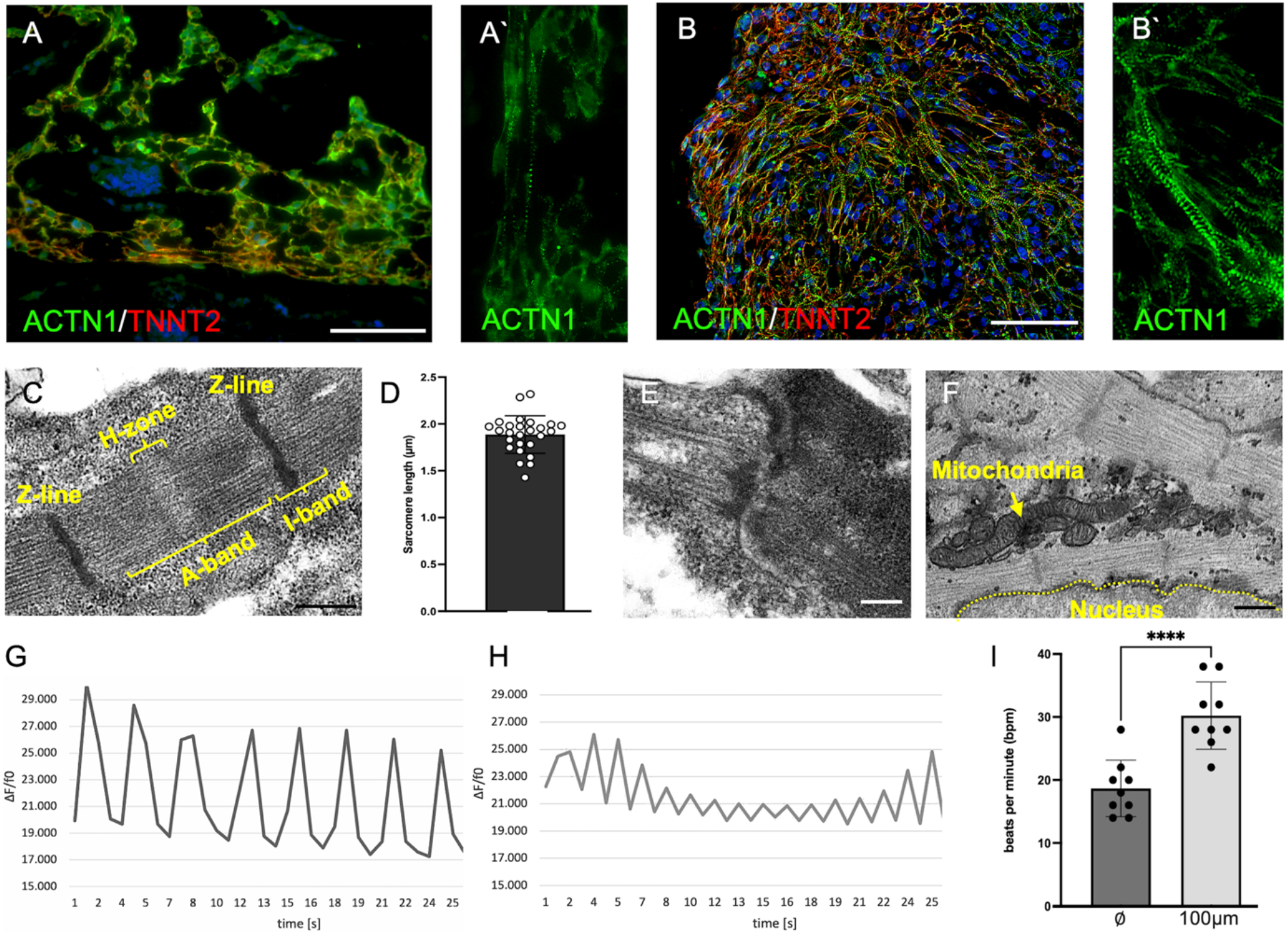
Cardiomyocyte Maturation in Organoids. **A)** Immature cardiomyocytes in day 8 organoids. Scale: 50 µm **A’)** Few sarcomere-like structures indicated by rarely occurring cross-striated staining pattern of ACTN1/TNNT2 with limited number of repeats. **B)** Cardiac organoids at day 35. Scale: 50 µm **B’)** Highly repetitive and well-developed cross-striated staining pattern of ACTN1 and TNNT2 indicating cardiomyocyte maturation. **C)** Sarcomeres at day 35 show a clear banding, with repetitive Z-lines, distinct I-bands, A-bands, and an H-zone. Scale: 500 nm **D)** Sarcomere length measurements determine a sarcomere length of 1.9 µm (± 0.2; n=26). **E)** Intercalated disc-like structures were observed. Scale: 500 nm **F)** Transmission electron microscopy reveals multiple mitochondria adjacent to sarcomeres. Scale: 200 nm. **G-I)** Beating frequency was measured in the absence and presence of 100µM Isoproterenol using calcium imaging. **G)** Calcium transients before stimulation. **H)** Calcium transients after stimulation with Isoproterenol. I) Cardiomyocytes increase their beating frequency from approximately 18 bpm (± 4.47) to 30 bpm (± 5.33) after treatment (n=9; P < 0.0001).

We also detected PECAM1^+^ endothelial cells in cardiac organoids. At day 8, PECAM1 staining reveals islands of roundish cells as well as primitive endothelial cords between patches of ACTN1^+^ cardiac tissue (Fig. 3A). At day 35, PECAM1^+^ cells form a network of endothelial cords throughout the cardiac tissue, and vessel-like structures with a clearly visible lumen are detected (Fig. 3B). Transmission electron microscopy confirms lumen formation and demonstrates the presence of larger and smaller vessels (Fig. 3C-D). Moreover, whole-mount immunofluorescent tissue clearing analysis shows the development of a continuous and branched vascular network (PECAM1^+^) within the cardiac muscle (ACTN1^+^) (Fig. 3E). Interestingly, PECAM1^+^ cell clusters are found within the lumen as well as the wall of larger endothelium-lined cavities. Besides PECAM1, the endothelial cells also express CD34 and some cells near the observed PECAM1^+^ clusters are CD45^+^, indicating hematopoiesis from hemogenic endothelium. (Fig. 3F-H). Intra-luminal hematopoietic clusters can be also visualized by electron microscopy (Fig. 3I), with cells showing a gradual condensation of the nucleus as observed during erythropoiesis *in vivo* (Fig. 3J). H&E staining reveals vessels filled with hematopoietic cells, reminiscent of vessels in the early embryo. (Fig. 3K, Video S1). We detect different stages of erythropoiesis (Fig. 3L) as well as biconvex, disc-shaped erythrocyte-like cells (Fig. 3M). Besides the intraluminal hematopoietic cells, AIF1^+^ macrophages are found near the PECAM1^+^ vessels (Fig. 3N). Macrophages are also present within the cardiac tissue (Fig. 3O), forming close contact sites with TNNT2^+^ cardiomyocytes (Fig. 3P).

**Figure 3.**
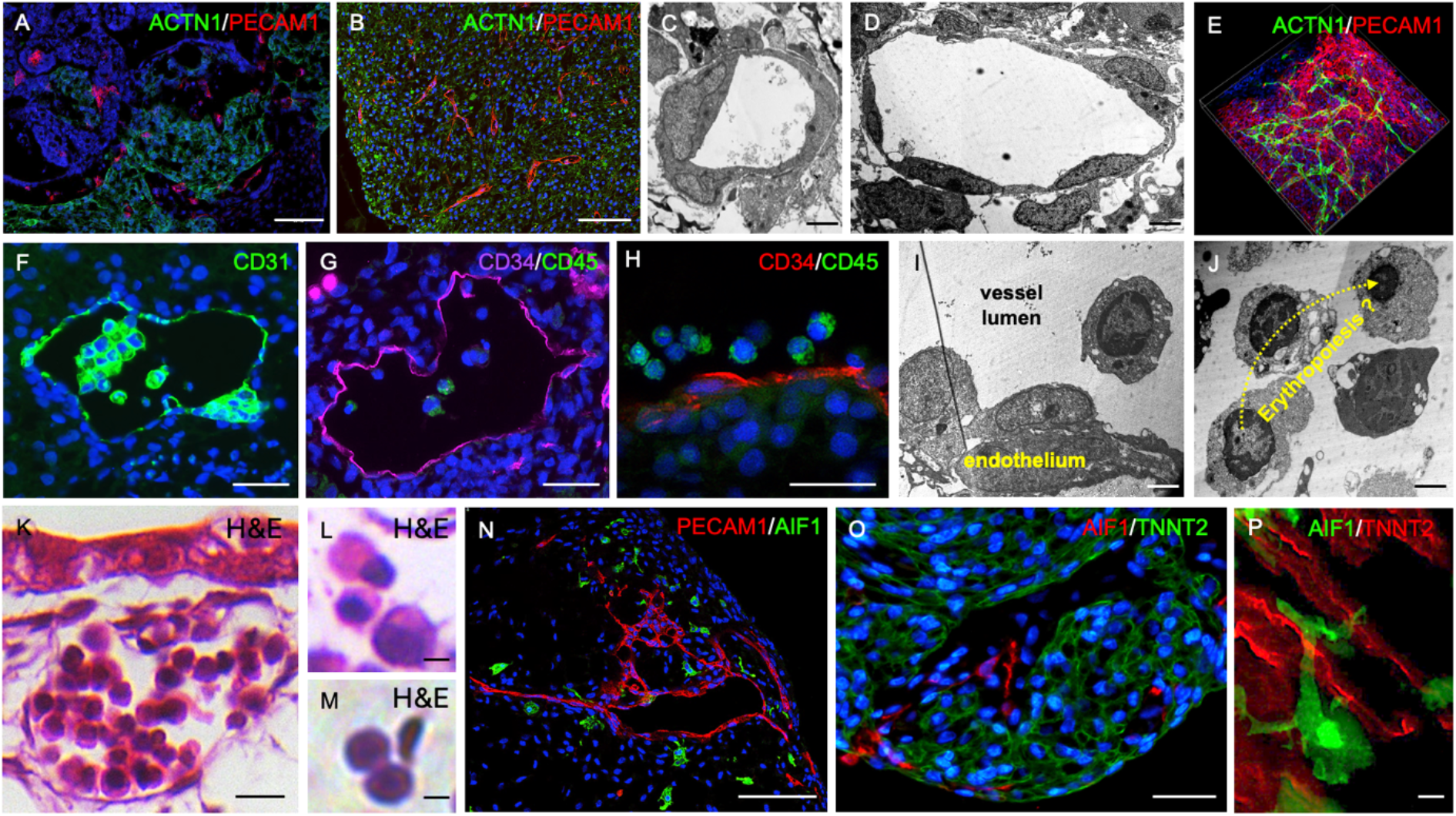
Vasculogenesis and Hematopoiesis in Cardiac Organoids. **A)** PECAM1^+^ cell islands and endothelial cords are detectable in day 8 organoids. Scale: 50 µm **B)** PECAM1^+^ cells form vessel-like structures within the cardiac tissue (ACTN1^+^) by day 35. Scale: 50 µm **C-D)** Transmission electron microscopy shows vessels with smaller (C) and larger (D) lumen diameter. Scale: 200 nm **E)** These vessels form a branched vascular network spanning the whole cardiac organoid, as seen in whole mount tissue clearing. **F)** PECAM1^+^ intraluminal clusters are detected within endothelium-lined vessels. Scale: 20 µm **G-H)** Intraluminal cells expressing CD34 and CD45. Scale: 20 µm **I)** Hemogenic endothelium-like structures detected by transmission electron microscopy. Scale: 100 nm **J)** Intraluminal cells undergo chromatin condensation reminiscent of erythropoiesis. Scale: 100 nm **K-M)** H&E staining of vessels with intraluminal hematopoietic cells. Different stages of erythropoiesis (L), as well as biconvex, disc-shaped erythrocyte-like cells (M) are detectable. Scale: 10 µm in K and 5 µM in L/M. **N)** AIF1^+^ macrophages are found near vessel-like structures. Scale: 50 µm **O)** Macrophages are also found within the musculature of the cardiac organoids. Scale: 20 µm **P)** Close interaction of AIF1^+^ macrophages with TNNT2^+^ cardiomyocytes revealed at high magnification. Scale: 5 µm.

We dissociated day 35 organoids and performed single-cell RNA sequencing to determine their exact cellular composition. The analyses reveal that organoids contain fibroblast-like cells, cardiomyocytes, endothelial cells, and macrophages (Fig. 4A). In addition, we find cells with an expression profile of intestinal epithelial cells, likely originating from the epithelial structures observed in the center of the organoid at day 8 (Fig. 1F). Each of the cell populations contain a subset of proliferative *MKI67*^+^ cells (Fig. 4B). Immunofluorescence analyses detecting MKI67 in TNNT2^+^ cardiac tissue as well as TNNT2^-^ non-myocytes confirm this result (Fig. 4C-D). Fig. 4E shows a heatmap of selected genes, which served as the basis for assigning the clusters to their cellular identities.

**Figure 4.**
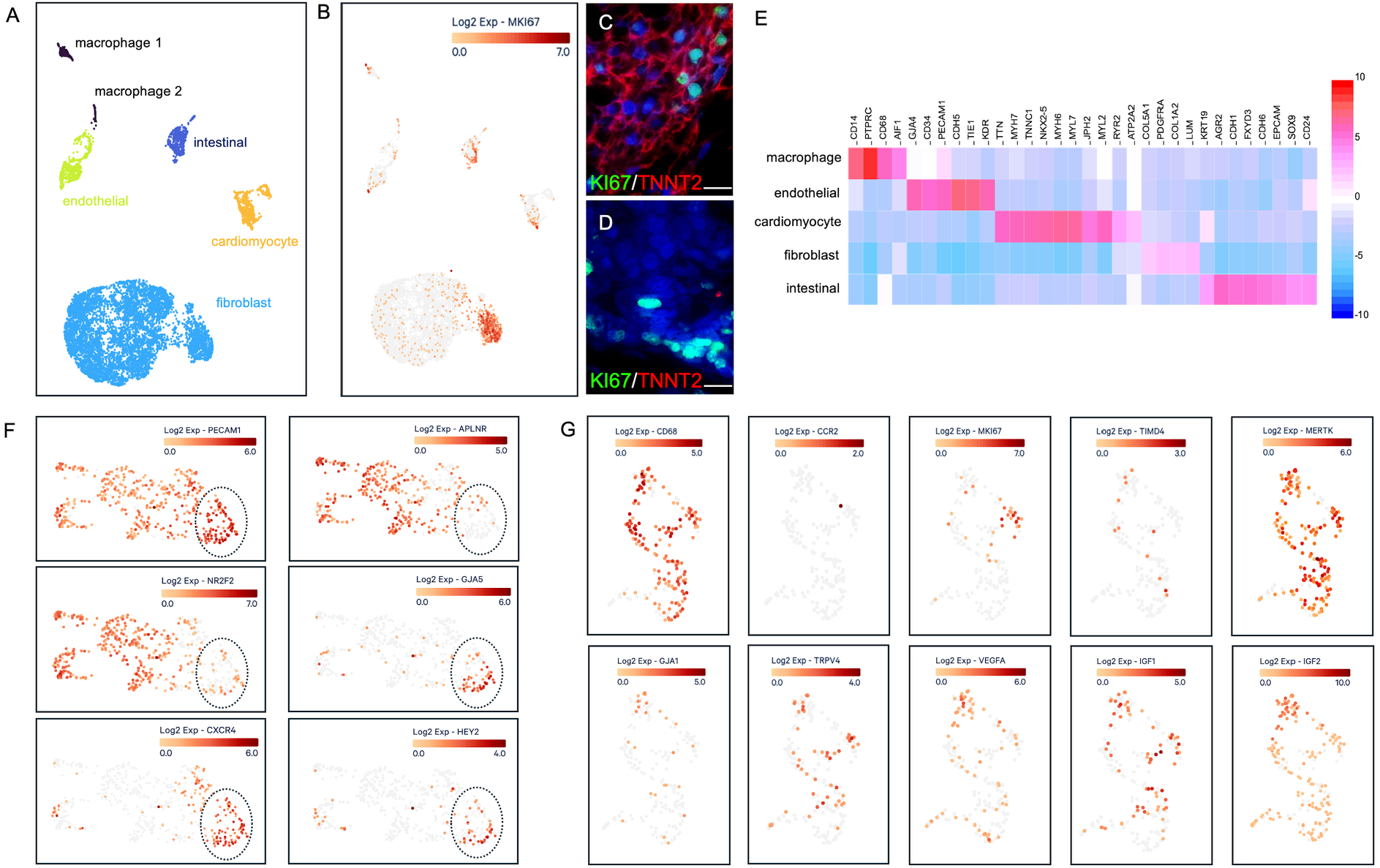
Single-cell RNA-sequencing analysis of cardiac organoids. **A)** Single-cell RNA-sequencing revealed cell clusters with characteristic gene expression profiles of fibroblasts, cardiomyocytes, endothelial cells, macrophages, and intestinal epithelial cells. **B)** Within all identified clusters, proliferating *MKI67*^*+*^ cells were observed. **C-D)** The scRNA-seq result was confirmed by immunofluorescence analyses detecting MKI67 in cardiomyocytes as well as non-cardiomyocyte tissue within the organoid. Scale: 10 µm. **E)** Heatmap of genes that were used as markers to identify the clusters depicted in A. **F)** Detailed analyses of scRNA-seq data of the endothelial cell cluster revealed cells with arterial identity (black dotted circle, GJA5^high^, CXCR4^high^, HEY2^high^, NR2F2^low^, APLNR^low^), supporting the assumption of hemogenic endothelium development. **G)** This hemogenic endothelium gives rise to *CCR2*^*-*^ tissue-resident macrophages that contain a subpopulation of proliferating cells (*MKI67*^*+*^) and express genes characteristic of tissue-resident cardiac macrophages such as *TIMD4, MERTK, GJA1*, and *TRPV4*, as well as genes for growth factors involved in cardiac development and remodeling (*VEGFA, IGF1, IGF2*).

Next, we investigated the endothelial cluster in more detail. All endothelial cells express *PECAM1*. However, a subset of cells (black dotted line) shows low expression of venous genes (*APLNR* and *NR2F2*) but high expression of arterial markers (*GJA5, CXCR4*, and *HEY2*) (Fig. 4F). Hemogenic endothelium is found in vessels with arterial identity, and therefore these cells could contribute to the production of hematopoietic cells in the cardiac organoids.

Finally, we examined the macrophage cell cluster (Fig. 4G). The cells express the macrophage marker *CD68* but are negative for *CCR2*, a hallmark of tissue-resident macrophages of embryonic origin (Bajpai et al., 2018). In the heart, a subset of CCR2-tissue-resident macrophages was described to be also TIMD4^+^ (Dick et al., 2019). Therefore, we checked for this marker and, indeed, a few *TIMD4*^*+*^ macrophages are detectable.

Resident macrophages contribute to cardiac function by MERTK dependent phagocytosis of exopheres, e.g. to remove non-functional mitochondria (Nicolas-Avila et al., 2020). We found that almost all cardioid macrophages express this receptor.

*In vivo*, cardiomyocytes and macrophages interact with each other through gap junctions made of GJA1 (Hulsmans et al., 2017). *GJA1* expression is detected in a subset of macrophages in the organoid. Moreover, TRPV4-expressing macrophages can be activated by mechanical stretch triggering the release of growth factors that contribute to cardiac remodeling (Wong et al., 2021). In our cultures, a subset of *CCR2-* macrophages express *TRPV4*. Additionally, they express mRNA coding the growth factors VEGFA, IGF1, and IGF2, which are important for cardiac remodeling (Fig. 4G). In summary, the scRNA sequencing results demonstrate that macrophages in organoids exhibit various features observed in cardiac tissue resident macrophages *in vivo*.

## Discussion

Here, we describe complex cardiac organoids generated from human iPSCs that consist of cardiomyocytes, fibroblasts, and endothelial cells. Cardiomyocytes undergo progressive maturation over time, developing more organized sarcomeres and an increase in spontaneous rhythmic contractions. More important, these cardiac organoids also contain a vascular network with artery-like endothelial domains that have blood-forming properties. A hemogenic endothelium gives rise to hematopoietic clusters inside the vessels, which generate erythrocytes and *CCR2-* tissue-resident macrophages (Bajpai et al., 2018). The macrophages infiltrate the heart muscle, coming into close contact with cardiomyocytes.

Our protocol is inspired by a previously published method (Lewis-Israeli et al., 2021). However, we replaced the WNT antagonist WNT-C59 by IWP2 and apply it at a low concentration of 2 µM. This leads to the appearance of macrophages and other hematopoietic cells which are not observed in the original protocol (Lewis-Israeli et al., 2021) (Volmert et al., 2023) (Kostina et al., 2023) (O’Hern et al., 2024). We hypothesize that the low dose of IWP2 (2 µM) leads to less tight and less efficient WNT inhibition, making the culture more permissive for the development of hematopoietic cells alongside the cardiomyocyte lineage. Notably, if IWP2 is used for cardiac organoid differentiation, it is usually applied at higher concentration of 5 µM (Drakhlis et al., 2021). IWP2 and WNT-C59 are both PORCN inhibitors (Shah et al., 2021), however, IWP2 was previously used for the specification of primitive hematopoietic cells from human iPSCs (Sturgeon et al., 2014) and was therefore favored.

Our lab previously demonstrated that vascularized mesodermal organoids give rise to microglia-like cells when co-cultured with neural organoids (Worsdorfer et al., 2019) demonstrating their hematopoietic potential. Recent, studies show that this hematopoietic capacity can be further enhanced by supplementing the cultures with hematopoietic growth factors such as SCF, IL-3, IL-6, TPO, and M-CSF, resulting in the formation of blood-producing bone marrow organoids (Frenz-Wiessner et al., 2024) (Olijnik et al., 2024). A recent study further showed that applying a precisely timed sequence of four complex growth factor cocktails to cardiac organoids can induce CD45^+^ hematopoietic cells (Dardano et al., 2024). Here, we show that hematopoiesis and macrophage formation can innately occur without adding additional growth factors, when low concentrations of IWP2 are used. Nevertheless, incorporating hematopoietic growth factors into our easy and cost-efficient protocol may offer opportunities for further optimization and will be tested in future experiments.

Recent studies suggest that macrophages form close connections with cardiomyocytes, either through focal adhesion complexes or gap junctions (Wong et al., 2021) (Hulsmans et al., 2017). Macrophages that connect to cardiomyocytes via focal adhesion complexes respond to mechanical stretch via a TRPV4-mediated signaling pathway, which controls growth factor expression and growth factor-induced cardiac remodeling. Gap junctions between macrophages and cardiomyocytes play a role in facilitating electrical conduction. Moreover, cardiac macrophages are closely associated with cardiomyocytes and help remove defective mitochondria through exosphere phagocytosis. Depletion of macrophages or defects in the phagocytic receptor MERTK lead to impaired autophagy, accumulation of defective mitochondria, and consequently, metabolic changes and heart dysfunction (Nicolas-Avila et al., 2020). All these findings show that tissue-resident macrophages play an important role in maintaining functional cardiac tissue and are probably also involved in cardiac development and tissue maturation. This assumption is supported by recent studies showing that adding macrophages to engineered heart tissue improves contractile strength, functionality, and long-term vascularization (Landau et al., 2024) (Lock et al., 2024) (Hamidzada et al., 2024).

Whether macrophages fulfill similar functions in the described cardiac organoids, leading to more mature and functional tissue models with properties like the heart *in vivo*, needs to be carefully investigated in future studies and is beyond the scope of this brief report. However, our single-cell RNA sequencing data indicate that the macrophages in the organoid display a tissue-resident macrophage-like expression profile and that genes involved in the processes described above, such as *TRPV4, CX43, MERTK* and *IGF1and VEGFA* are expressed.

In conclusion, we present a novel cardiac organoid model containing tissue-resident macrophages, an important cell type of the heart tissue that has so far been missing in cardiac organoids.

## Materials and Methods

### Human iPSC culture

All experiments were conducted using the human induced pluripotent stem cell (iPSC) line NHDF iPSC. The reproducibility of cardiac organoid generation and innate macrophage development was confirmed using an independent iPSC line, KK iPSC. NHDF iPSCs were generated from commercially available male juvenile human dermal fibroblasts (juvenile NHDF, C-12300, PromoCell) via reprogramming with the hSTEMCCA lentiviral construct (Sommer et al., 2012). KK iPSCs were derived from dermal fibroblasts isolated from a postmortem skin biopsy of a 94-year-old female donor who had consented to body donation through our anatomical institute. These fibroblasts were reprogrammed using a commercially available Sendai virus-based reprogramming kit (CytoTune 2.0 Sendai Reprogramming Vectors, Ref# A16517; Lot# A16517, Invitrogen).

Human iPSCs were cultured on Matrigel-coated 6-well plates in StemMACS iPS-Brew medium (Miltenyi). Cells were passaged upon reaching approximately 80% confluency using Accutase. Following passaging, cells were reseeded at the desired density in StemMACS iPS-Brew medium supplemented with 10 µM ROCK inhibitor Y-27632.

### Generation of cardiac organoids

To generate cardiac organoids, a single-cell suspension of iPSCs was prepared using Accutase, and 10,000 cells were seeded per well into a round-bottom, ultra-low attachment 96-well plate in 100 µL of StemMACS iPS-Brew medium (Miltenyi), supplemented with 10 µM ROCK inhibitor Y-27632 (designated as day -1). The plate was incubated in a humidified incubator at 37 °C with 5% CO_2_ on an orbital shaker for 24 hours. By day 0, single iPSC aggregates had formed in each well.

To initiate differentiation, 20 µL of medium was removed from each well and replaced with 166.6 µL of mesoderm induction medium (MIM), consisting of RPMI 1640 (Gibco, Cat# 11875-093) supplemented with 2× B27 without insulin (Life Technologies), 6 µM CHIR99021 (Sigma-Aldrich), 1.875 ng/mL BMP4 (PreproTech), and 1 ng/mL Activin A (Miltenyi Biotech).

In the following steps, 2/3 (166.5 µl) of the medium are removed per medium change and replaced by new medium.

After 24 hours (day 1), the medium was changed to Basic Medium 1 (BM1), composed of RPMI 1640 with 2x B27 without insulin, and incubated for another 24 hours. On day 2, to promote cardiac mesoderm specification, the medium was replaced with Cardiac Specification Medium (CSM), consisting of RPMI 1640 supplemented with 2% B27 without insulin and 3 µM IWP2 (Miltenyi Biotech). Aggregates were maintained in CSM for 48 hours (until day 4), followed by a medium change back to BM1 for an additional 48 hours.

From day 6 onward, for cardiac maturation and maintenance, the medium was switched to Basic Medium 2 (BM2), composed of RPMI 1640 supplemented with 2% B27 (Life Technologies). BM2 was refreshed every other day until the organoids were either fixed for histological sectioning, dissociated for single-cell RNA sequencing, or subjected to calcium imaging or other analyses.

A schematic overview of the protocol timeline is provided in Figure 1A.

### Dissociation of Cardioids

Cardiac organoids were dissociated using a cold-activated protease method. The dissociation solution consisted of 10 mg/mL Bacillus licheniformis protease (Creative Enzymes, NATE-0633) and 125 U/mL DNase I (Merck, DN25) in ice-cold DPBS (Sigma-Aldrich) supplemented with 5 mM CaCl2 (Carl Roth), prepared in a total volume of 1 mL.

Organoids were washed once with ice-cold PBS containing 10% fetal calf serum (FCS) (Biowest). After discarding the washing solution, 1 mL of the dissociation solution was added. Organoids were incubated on ice for 15 minutes, during which they were gently triturated every minute using a P1000 pipette.

The dissociation reaction was stopped by adding 1 mL of PBS with 10% FCS, followed by centrifugation at 300 x g for 3 minutes. The resulting cell pellet was resuspended in 10 mL of PBS with 10% FCS and filtered through a pre-wetted 40 µm cell strainer.

Single cells were counted using a Neubauer hemocytometer, then centrifuged again at 300 x g for 3 minutes. The supernatant was discarded, and the cell pellet was resuspended in PBS containing 0.04% bovine serum albumin (BSA) for scRNA sequencing analysis.

### Single-cell RNA-sequencing

Library preparation was performed according to the manufacturer’s instructions (Chromium Next GEM Single Cell 3’ v3.1 protocol CG000390 Rev B, 10x Genomics). Briefly, cells were resuspended in the master mix and loaded, along with partitioning oil and gel beads, into the microfluidic chip to generate a gel beademulsion (GEM). The poly-A RNA from the cell lysate contained within each droplet was reverse transcribed into cDNA, which included an Illumina R1 primer sequence, a Unique Molecular Identifier (UMI), and the 10x Barcode.

The pooled, barcoded cDNA was then purified using Silane DynaBeads, amplified by PCR, and size-selected using SPRIselect reagent to isolate appropriately sized fragments for subsequent library construction. During library construction, the Illumina R2 primer sequence, paired-end constructs with P5 and P7 sequences, and a sample index were added.

Library quantification and quality control were performed using a Qubit™ 4.0 Fluorometer (Thermo Fisher) and a 2100 Bioanalyzer with the High Sensitivity DNA Kit (Agilent). Sequencing was carried out on a NovaSeq 6000 sequencer (Illumina) using an S2 flow cell.

Downstream data analysis was conducted using the Loupe Browser v8 (10x Genomics).

### Immunofluorescence Analyses

Cardiac organoids were fixed in 4% paraformaldehyde (PFA; Sigma-Aldrich) at 4 °C overnight, washed in phosphate-buffered saline (PBS; Sigma-Aldrich), and embedded in 1% agarose gel (Biozym). Subsequently, 5 µm paraffin sections were prepared.

For immunofluorescence staining, sections were deparaffinized, rehydrated, and subjected to antigen retrieval using 10 mM sodium citrate buffer (pH 6.0). Primary antibodies against PECAM1 (CD31) (DAKO, M0823), Vimentin (Invitrogen, MA5-6409), ACTN1 (Abcam, Ab68167), KDR (Miltenyi, 130-125-988), TBXT (R&D Systems, AF2085), TNNT2 (Life Technologies, MA512960), CD34 (Life Technologies, MA5-32059), AIF1 (WAKO/Fuji Film, 019-19741) and CD45 (Life Technologies, 14-0451-82) were diluted in NBS blocking solution and incubated overnight at 4 °C. Secondary antibodies conjugated to Cy2, Cy3, or Cy5 fluorophores (Dianova) were diluted in PBS and applied for 1 hour at room temperature. Nuclei were counterstained with DAPI (Roche).

Additionally, sections were stained with eosin (AppliChem) and hematoxylin (Chroma) for histological evaluation. Images were acquired using an Axiovert 40 CFL microscope (Carl Zeiss Microscopy GmbH), a Keyence BZ-X fluorescence microscope, and a Nikon Eclipse Ti confocal laser scanning microscope.

### Tissue Clearing

Tissue clearing was performed according to a previously published protocol (Worsdorfer et al., 2020). For immunofluorescence staining of cleared cardiac organoids, primary antibodies against PECAM1 (DAKO, M0823), ACTN1 (Abcam, Ab68167), TNNT2 (Life Technologies, MA512960), and AIF1 (WAKO/Fuji Film, 019-19741) were used.

Imaging was carried out using a Nikon Eclipse Ti confocal laser scanning microscope equipped with long working distance air objectives (4x and 20x) to acquire z-stack images. Three-dimensional reconstruction of the acquired image stacks was performed using Fiji (ImageJ) or Nikon NIS-Elements Confocal software.

### Transmission Electron Microscopy

Cardiac organoids were washed with PBS and kept on ice for 15 minutes prior to fixation. Organoids were fixed overnight at 4 °C in a solution containing 2.5% glutaraldehyde, 4% paraformaldehyde (PFA), and 2 mM CaCl_2_ in 0.1 M cacodylate buffer (composed of 50 mM cacodylate, 50 mM KCl, and 2.5 mM MgCl_2_, pH 7.2). Following fixation, samples were washed with 0.1 M cacodylate buffer and subjected to a second fixation step using 1% osmium tetroxide in 0.1 M cacodylate buffer for 1 hour.

After osmium fixation, organoids were washed for 10 minutes in 0.1 M cacodylate buffer, followed by two 10-minute washes in double-distilled water (ddH_2_O). Samples were then incubated in 8% uranyl acetate substitute for 1 hour and washed again twice for 10 minutes in ddH_2_O.

Dehydration was carried out using a graded ethanol series (30%, 50%, 70%, 90%, and two changes of 100% ethanol), with each step lasting 10 minutes. Organoids were then incubated in propylene oxide (PO) for two 30-minute intervals, followed by overnight incubation in a 1:1 mixture of PO and Epon 812 resin.

The following day, samples were incubated in pure Epon for 2 hours and embedded by polymerizing the resin at 60 °C for 48 hours. Ultrathin sections were prepared using an ultramicrotome, collected on nickel grids, and post-stained with 2.5% uranyl acetate and 0.2% lead citrate. Finally, specimens were analyzed using a LEO AB 912 transmission electron microscope (Zeiss).

### Semiquantitative RT Polymerase Chain Reaction

For polymerase chain reaction (PCR), samples were disrupted using ultrasonic sonication (Ultrasound Processor, 20 kHz, 80 W; power setting 30, 3 × 20 s pulses with 3 s intervals), followed by RNA extraction using the Direct-zol RNA MiniPrep Plus Kit (Zymo Research). cDNA synthesis was performed using the GoScript™ Reverse Transcriptase (Promega), following the manufacturer’s instructions. PCR amplification was carried out using the Red MasterMix (2×) Taq PCR MasterMix (Genaxxon). The following primer pairs were used:

GAPDH

Forward:TGACAACTTTGGTATCGTGGA

Reverse: CCAGTAGAGGCAGGGATGAT

NKX2.5

Forward:CCAAGGACCCTAGAGCCGAA

Reverse: ATAGGCGGGGTAGGCGTTAT

TNNT2

Forward:GGAGGAGTCCAAACCAAAGCC

Reverse: TCAAAGTCCACTCTCTCTCCATC

DES

Forward:TCGGCTCTAAGGGCTCCTC

Reverse: CGTGGTCAGAAACTCCTGGTT

ISL1

Forward:GCGGAGTGTAATCAGTATTTGGA

Reverse: GCATTTGATCCCGTACAACCT

KDR

Forward:GGCCCAATAATCAGAGTGGCA

Reverse: CCAGTGTCATTTCCGATCACTTT

PECAM1

Forward:AACAGTGTTGACATGAAGAGCC

Reverse: TGTAAAACAGCACGTCATCCTT

TBXT

Forward:TATGAGCCTCGAATCCACATAGT

Reverse: CCTCGTTCTGATAAGCAGTCAC

GATA4

Forward:CGACACCCCAATCTCGATATG

Reverse: GTTGCACAGATAGTGACCCGT

TBX5

Forward:CTGTGGCTAAAATTCCACGAAGT

Reverse: GTGATCGTCGGCAGGTACAAT

RYR2

Forward:ACAACAGAAGCTATGCTTGGC

Reverse: GAGGAGTGTTCGATGACCACC

### Calcium Imaging

For calcium imaging, cardioids were incubated with 1 µM Fluo-4 AM (Gibco) in DMEM (Gibco) at 37 °C for 30 minutes. For stimulation experiments, the Fluo-4 AM-containing medium was replaced with DMEM supplemented with 100 µM isoproterenol (Sigma, dissolved in ethanol) and incubated at 37 °C for an additional 30 minutes, followed by a final medium change.

Dynamic fluorescence changes were recorded before and after treatment using a Leica DM IL LED microscope equipped with a Leica EL6000 external fluorescence light source and a Leica DFC450C camera. Videos were processed using Leica Application Suite Version 4.12.0. Fluorescence data were analyzed using Fiji (ImageJ) and Microsoft Excel.

### Statistics

All statistical analyses were performed using GraphPad Prism version 10. Data are presented as bar graphs showing mean ± standard error of the mean (SEM), with asterisks indicating statistical significance where applicable.

Normality of data distribution was assessed using the Shapiro–Wilk test. For normally distributed data, statistical significance was determined using one-way ANOVA followed by Bonferroni’s multiple comparison test.

## Acknowledgement

We thank Doris Dettelbacher-Weber, Martina Gebhardt, Erna Kleinschroth, Karin Reinfurt-Gehm, Ursula Roth, Sieglinde Schenk, Elke Varin, and Lisa Wittstatt for their excellent technical assistance, as well as all members of the Stem Cell and Regenerative Medicine Group for their support and valuable discussions. We thank Fabian Imdahl and the Core Unit Systems Medicine at the University of Würzburg for excellent technical support and RNA-seq data generation. This work was supported by the Interdisciplinary Center for Clinical Research (IZKF) Würzburg (project Z-06).This work was further supported by grants from the IZKF Würzburg (project E-D-410) to P.W. and B.G., and by the German Research Foundation (DFG) through the Collaborative Research Center TRR225-B04 to S.E. This publication was supported by the Open Access Publication Fund of the University of Würzburg. Figure 1A was created using BioRender (BioRender.com).

## Notes

### Competing Interest Statement

The authors have declared no competing interest.

